# Short-term audiovisual spatial training enhances electrophysiological correlates of auditory selective spatial attention

**DOI:** 10.1101/2020.12.23.424131

**Authors:** Christina Hanenberg, Michael-Christian Schlüter, Stephan Getzmann, Jörg Lewald

**Affiliations:** Ruhr University Bochum, Faculty of Psychology, D-44780 Bochum, Germany; Leibniz Research Centre for Working Environment and Human Factors, D-44139 Dortmund, Germany

## Abstract

Audiovisual cross-modal training has been proposed as a tool to improve human spatial hearing. Here, we investigated training-induced modulations of auditory-evoked event-related potential (ERP) components that have been associated with processes of auditory selective spatial attention when a speaker of interest has to be localized in a multiple speaker (“cocktail-party”) scenario. Forty-five healthy subjects were tested, including younger (19-29 yrs; *n* = 21) and older (66-76 yrs; *n* = 24) age groups. Three conditions of short-term training (duration 15 minutes) were compared, requiring localization of non-speech targets under “cocktail-party” conditions with either (1) synchronous presentation of co-localized auditory-target and visual stimuli (audiovisual-congruency training) or (2) immediate visual feedback on correct or incorrect localization responses (visual-feedback training), or (3) presentation of spatially incongruent auditory-target and visual stimuli presented at random positions with synchronous onset (control condition). Prior to and after training, subjects were tested in an auditory spatial attention task (15 minutes), requiring localization of a predefined spoken word out of three distractor words, which were presented with synchronous stimulus onset from different positions. Peaks of ERP components were analyzed with a specific focus on the N2, which is known to be a correlate of auditory selective spatial attention. N2 amplitudes were significantly larger after audiovisual-congruency training compared with the remaining training conditions for younger, but not older, subjects. Also, at the time of the N2, electrical imaging revealed an enhancement of electrical activity induced by audiovisual-congruency training in dorsolateral prefrontal cortex (Brodmann area 9) for the younger group. These findings suggest that cross-modal processes induced by audiovisual-congruency training under “cocktail-party” conditions at a short time scale resulted in an enhancement of correlates of auditory selective spatial attention.

## Introduction

Numerous lines of human and animal research have provided clear evidence that the representation of sound sources in space can be modulated by vision. In particular, in the so-called ventriloquism effect the perceived sound location is shifted toward a spatially disparate, temporally coincident visual event (e.g., Klemm, 1910; Thomas, 1941; Jackson, 1953; Howard & Templeton, 1966; Jack & Thurlow, 1973; Lewald et al., 2001; Lewald & Guski, 2003; for review, see Radeau, 1994). Moreover, exposure to a consistent audiovisual spatial disparity over a certain period of time can induce a systematic shift in sound localization such that the representation of the auditory space is shifted to that of the visual space (Helmholtz, 1867; Stratton, 1896, 1897; Held 1955; Kalil & Freedman 1967; Canon 1970, 1971; Radeau & Bertelson, 1977, 1978; Recanzone, 1998; Lewald, 2002b). These cross-modal adaptive changes, which can emerge over short time scales from seconds to minutes (cf. Bosen et al., 2018), have been termed ventriloquism after-effect. Inspired by animal studies, which demonstrated similar (though more long-term) plasticity of auditory and visual neural representations (Knudsen & Knudsen, 1985, 1989; Brainard & Knudsen, 1993; Knudsen, 1999; Hyde & Knudsen, 2000, 2002; Zheng & Knudsen, 2001), experiments on this phenomenon led to the conception that vision calibrates human auditory spatial perception (Recanzone, 1998; Lewald, 2002b). Results obtained in blind and blindfolded sighted humans as well as in patients with visual-field loss demonstrating specific alterations of sound localization were in accordance with this view (e.g., Zwiers et al., 2001a, b; Lewald, 2002a, c, 2013; Lewald et al., 2009a, b, 2013; Feierabend et al., 2019).

On the basis of neuroscience studies on multisensory learning (for review, see Shams & Seitz, 2008), approaches of sensory training have been developed, in which auditory stimuli were presented in spatio-temporal alignment with visual stimuli. For example, in patients with pure hemianopia, who suffer from a loss of one half of the visual field due to brain damage while having sufficient audiospatial performance (Lewald et al., 2009b, 2013), neuro-rehabilitative audiovisual training or even auditory unimodal stimulation have been shown to induce long-lasting improvements of visual functions in the anopic hemifield (Bolognini et al., 2005; Passamonti et al., 2009; Lewald et al., 2012). Conversely, cross-modal approaches of sensory training have also demonstrated improvements of auditory functions, suggesting that persons with hearing impairments could benefit from them (Zahorik et al., 2006; Strelnikov et al., 2011; Majdak et al., 2013; Kuk et al., 2014; Grasso et al., 2016; Cai et al., 2018). In the audiospatial domain, audiovisual training has been shown to significantly increase the accuracy of localization of single sound sources under monaural (Strelnikov et al., 2011) and binaural conditions in healthy adults (Cai et al., 2018). In particular, auditory-visual training was found to induce a stronger improvement in sound localization compared to auditory-only training and a significant reduction of front-back confusion for both, trained and untrained sound positions (Cai et al., 2018).

While previous approaches to improve audiospatial performance by sensory training used single sound sources presented in isolation (e.g., Cai et al., 2018), spatial hearing in everyday life requires more complex functions of selective spatial attention, since auditory objects of interest have to be detected and localized among several distractor sound sources. Listening in such a “cocktail-party” situation (Cherry, 1953) is a remarkable ability of the human auditory system, which allows to orient the focus of attention to a sound source of interest in noisy, multi-speaker scenarios (for review, see Bregman, 1990; Bronkhorst, 2015). Such conditions of listening can be challenging already for normal-hearing people, but become substantially more difficult in hearing-impaired persons and at older age, resulting in serious restrictions of communication and social interaction in everyday life (Lewald & Hausmann, 2013; Getzmann et al., 2014, Getzmann, Wascher, et al., 2015; see also Pichora-Fuller et al., 2017). This leads to the question of whether hearing performance under these conditions could be improved by training interventions. On the basis of the previous sensory-training approaches mentioned above, it seems reasonable to assume that audiovisual stimulation could be an effective tool in this respect. Thus, in the present study a bimodal spatial training was developed in order to enhance brain functions associated with selective spatial attention under multiple-speaker conditions. Two types of sensory training were employed and compared with a control condition: *(1)* an audiovisual-congruency training, in which auditory targets were presented in spatiotemporal alignment with light stimuli, and *(2)* a visual-feedback training, in which correct responses on target location were indicated by light flashes. We hypothesized that audiovisual-congruency training may result in more effective learning due to the specific enhancement of multimodal brain circuits by audiovisual spatiotemporal alignment (Shams & Seitz, 2008).

The study was focused on training-induced modulations of event-related potential (ERP) components that have been associated with processes of auditory selective spatial attention in “cocktail-party” situations, using a multiple-speaker sound-localization task (Lewald et al., 2016; Hanenberg et al., 2019). In particular, we expected a training-induced increase of the N2 component. Localization of predefined auditory target stimuli in multiple-distracter environments has been shown to result in a substantially stronger N2 component of the ERP compared with single-source localization (Lewald & Getzmann, 2015). More generally, the N2 component has been regarded as a neural correlate of processes of cognitive control and orienting attention (Folstein & Van Petten, 2008; Potts, 2004), and has been related to conflict processing and suppression of irrelevant information in the auditory domain (Bertoli et al., 2005; Falkenstein et al., 2002; Getzmann, Wascher, et al., 2015). The N2 is usually reduced in older, compared with younger, adults (e.g., Anderer et al., 1996; Wascher & Getzmann, 2014). This has been interpreted to reflect an age-related less efficient inhibitory control over concurrent speech information (Getzmann, Wascher, et al., 2015; Getzmann, Hanenberg, et al., 2015), in line with the more general inhibitory deficit hypothesis (Hasher & Zacks, 1988; for review, see Gazzaley & D’Esposito, 2007). Therefore, as we were interested in potential age-related differences in training effects on ERPs (as demonstrated for the P300; e.g., Yang et al., 2018), groups of younger (19-29 yrs) and older participants (66-76 yrs) were compared.

## Materials and Methods

### Subjects

Forty-five adult subjects participated in this study, assigned to either a younger (*n* = 21; 12 women; mean age 25.0 yrs, SE 0.7 yrs, age range 19-29 yrs) or older group (*n* = 24; 12 women; mean age 71.0 yrs, SE 0.7 yrs, age range 66-76 yrs). Data from eight further participants (three younger and five older) were excluded from the analyses since the subjects responded in less than 50% of all trials. All subjects spoke German fluently and wrote with their right hand. Audiometric thresholds of all subjects were normal (mean across 11 pure-tone frequencies ≤ 25 dB hearing level for the younger group, and ≤ 40 dB hearing level for the older group; 0.125-8 kHz; Oscilla USB100, Inmedico, Lystrup, Denmark). This study conformed to the Code of Ethics of the World Medical Association (Declaration of Helsinki), printed in the British Medical Journal (18 July 1964) and was approved by the Ethical Committee of the Leibniz Research Centre for Working Environment and Human Factors, Dortmund. All subjects gave their written informed consent for participation.

### Experimental Setup

As in previous studies, the experiments were carried out in a dimly illuminated soundproof and echo-reduced room (5.0 × 3.3 × 2.4 m^3^; Getzmann et al., 2014). Subjects were seated on a comfortable chair, positioned with equal distances to the front-and side-walls of the room. At a distance of 1.5 m from the subject’s head, a semicircular array with four active broadband loudspeakers (SC5.9; Visaton, Haan, Germany; housing volume 340 cm^3^) was arranged in the horizontal plane at −60° and −20° to the left, and at 20° and 60° to the right of the subject’s median plane (Fig. 1A). The subject’s head position was stabilized by a chin rest. In the median plane of the subject’s head, a red light-emitting diode (LED; diameter 3 mm, luminous intensity 0.025 mcd), mounted at the central position of the semicircular array, served as fixation target during testing, while no fixation was required during training. For audiovisual-congruency and visual-feedback training (see *General Procedure),* an array consisting of a red (4 × 7 cm^2^; 70 cd/m^2^) and a white LED screen (4 × 7 cm^2^; 200 cd/m^2^) was mounted below each loudspeaker. The two screens were mounted below each other, with the white screen in the upper position, immediately adjacent to the lower edge of the loudspeaker housing (see Fig. 1B).

**Fig. 1.**
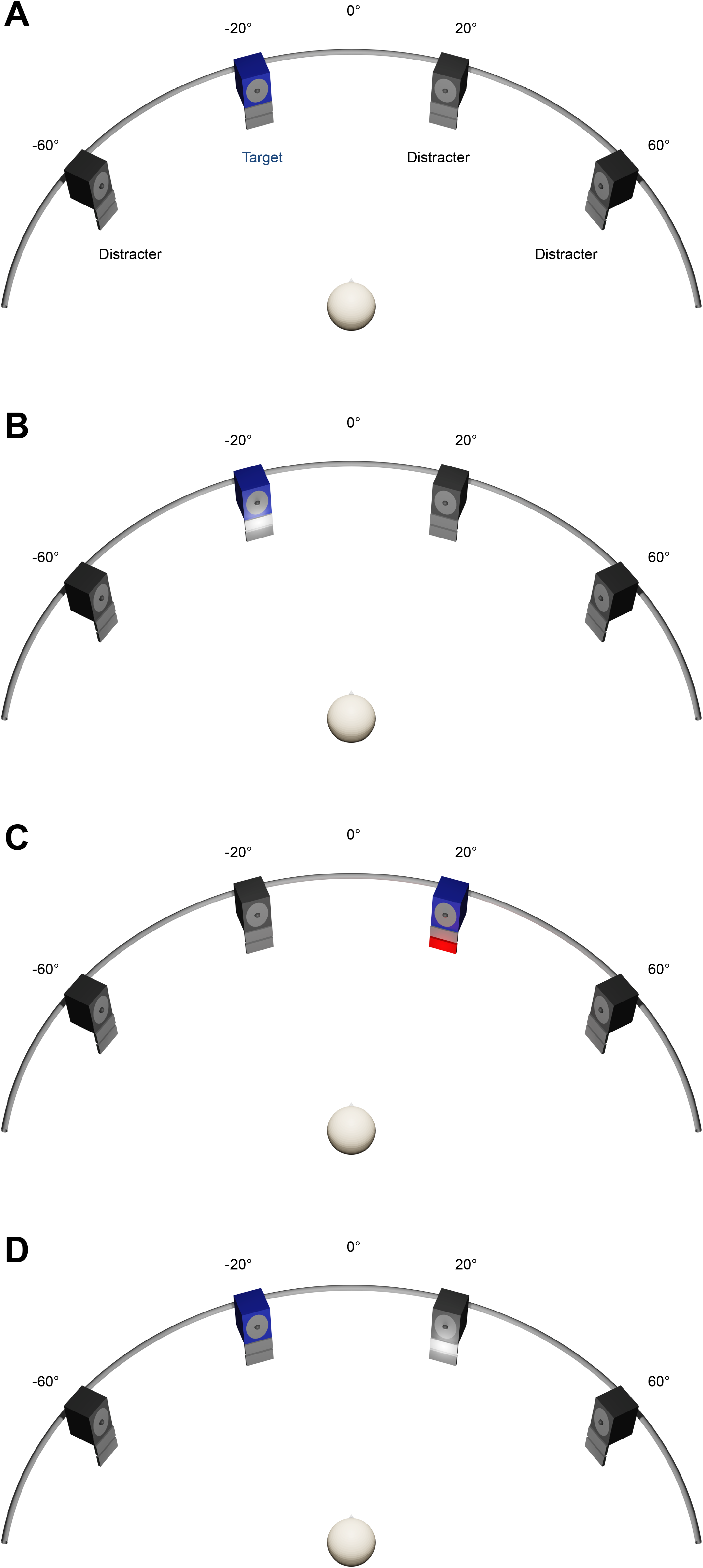
Experimental set-up. A: Auditory spatial attention task. Four numerals were presented simultaneously from four loudspeakers located at −60° and −20° to the left and 20° and 60° to the right. Subjects had to localize a predefined target numeral by using a four-button response box. B-D: Training Interventions. Four animal vocalizations were presented simultaneously from four loudspeakers located at −60°, −20°, 20°, and 60°. Subjects had to localize a predefined target vocalization by using the same response box as used in spatial attention task. In the audiovisual-congruency training (B), auditory target and visual stimuli were presented in spatiotemporal alignment. In the visual-feedback training (C), visual feedback on correct or incorrect localization was given immediately after each response. In the control condition (D), spatially incongruent auditory target and visual stimuli were presented at random positions with synchronous onset.

### Auditory Stimuli

The auditory stimuli used for testing were as in previous studies (for further details, see Lewald et al., 2016). Four different German one-syllable numerals, spoken by four different speakers (duration about 600 ms) were presented with synchronous stimulus onset from the four loudspeaker positions. The numerals used were “eins” (1), “vier” (4), “acht” (8), and “zehn” (10), each of them spoken by two male and two female native German speakers. Each numeral was presented from each of the four loudspeakers with equal frequency of occurrence at an overall sound pressure level of 66 dB(A), as was measured at the position of the subject’s head, using a sound level meter with a ½” free-field measuring microphone (Types 2226 and 4175, Brüel & Kjær, Nærum, Denmark).

Stimuli used for training have already been used previously (for details, see Lewald, 2016). Sounds consisted of animal vocalizations taken from a sound library (Marcell et al., 2000). Four different vocalizations (denoted ‘birds chirping’; ‘dog barking’; ‘frog’; ‘sheep’ in Marcell et al., 2000) were presented with synchronous stimulus onset from the four loudspeakers. Stimulus durations were adjusted to about 600 ms by cutting the original sound files. Stimuli were digitized at 48 kHz sampling rate and 16-bit resolution and converted to analogue form via a PC-controlled soundcard (Terrasoniq TS88 PCI, TerraTec Electronic, Nettetal, Germany). Each animal vocalization was presented from each of the four loudspeakers with equal frequency of occurrence. Stimuli were presented at an overall sound pressure level of 70 dB(A), measured as for the stimuli used for testing (see above).

### General Procedure

Following a within-subject repeated-measures crossover design, each subject was tested in three sessions conducted on different days, with intervals of at least one week and maximally three weeks in between. Each session included a different condition of training: a task of localization of a pre-defined target sound (animal vocalization) among 3 distracters was combined with *(1)* synchronous presentation of co-localized auditory and visual stimuli *(audiovisual-congruency), (2)* immediate visual feedback on correct or incorrect localization responses *(visual-feedback),* or *(3)* presentation of spatially incongruent auditory-target and visual stimuli presented at random positions with synchronous onset *(control condition).* The sequence of conditions was counterbalanced across subjects.

Each session comprised four blocks, three identical blocks of testing with an auditory spatial attention task (see below), each consisting of 288 trials (duration 15 min), and one training block (288-316 trials, duration 15.0-16.5 min). The first test block (pre-training) was used as baseline measurement, immediately followed by the training block. The second test block (post-training) was started immediately following the training, and the third test block (1-h post-training) began 60 min after the end of the training block. Between pre-training and training-blocks, the instructions given prior to experiments related to the specific training condition (see *Training Conditions)* were briefly repeated. Between post-training and 1-h post-training blocks, subjects were allowed to rest, remaining seated in the experimental chair. Prior to the experiment, all subjects were informed that they would receive a type of audiovisual training in each session.

### Auditory Spatial Attention Task

The auditory spatial attention task used for testing was similar to previous studies (Lewald et al., 2016; Hanenberg et al., 2019). In each trial, subjects had to localize a predefined target numeral out of three distracter numerals by pressing one out of four response buttons (Fig. 1A). The buttons were semi-circularly arranged on a response box, representing the four possible target positions (i.e., far left, mid left, mid right, far right). Subjects were instructed to use the right index finger for responding. For each subject, one numeral (1, 4, 8, or 10) was defined as target, which was kept constant for all measurements. Targets were counterbalanced across subjects. During testing, subjects had to to keep the eyes open and to fixate on the central LED to reduce artefacts due to eye movements and alpha activity in the EEG. Subjects were instructed to respond as accurately as possible within about 2 s after stimulus offset. Each trial lasted 3.125 s. Target position, distracter positions, and speakers changed between trials following a pseudo-random order (for details, see Lewald et al., 2016). The timing of the stimuli and the recording of the subjects’ responses were controlled by custom-written software. Before the beginning of the experiment, subjects completed about ten practice trials. They did not receive any feedback on their performance in the auditory spatial attention task during the experiment.

### Training Conditions

In the *audiovisual-congruency training* condition (Fig. 1B), the target sound appeared simultaneously with illumination of the white LED screen (duration 600 ms) mounted immediately below the target loudspeaker in 288 of a total of 316 trials (trial duration 3.125 s). To keep constant the subjects’ spatial attention, 28 of the trials were catch trials, in which target and LED screen appeared at incongruent positions. Subjects had to indicate sound locations using the response box, as described for the auditory spatial attention task (see above). Target sounds during training were assigned to target numerals during test blocks, that is, when the target numeral was ‘1’ during testing, the target animal vocalization during training was always ‘birds chirping’, ‘4’ was combined with ‘dog barking’, ‘8’ with ‘frog’, and ‘10’ with ‘sheep’. Subjects were informed that white LED screens did not reliably predict the position of the target and were instructed to rely on audition, not vision. Incorrect responses were indicated by flashing of the red LED screen (duration 600 ms) at the actual target position immediately after button pressing. Subjects were briefed about this procedure prior to training.

In the *visual-feedback training* condition (Fig. 1C), the sound-localization task and the presentation of auditory stimuli were as in the audiovisual-congruency training condition, except that catch trials were missing, thus resulting in a total of 288 trials. For visual feedback, each response was immediately followed by flashing of one of the two LED screens mounted below the loudspeaker that emitted the target sound (duration 600 ms). Correct responses were indicated by white light, incorrect responses by red light at the actual target position. Prior to training, the subject was informed about the visual feedback.

The *control* condition (Fig. 1D) was similar to the audiovisual-congruency training condition (288 trials). However, the white LED screen appeared at random positions, always diverging from the auditory target (duration 600 ms). Subjects were instructed to localize the predefined target sound while ignoring the light flashes. In this condition, no feedback was provided after pressing a button.

### Analysis of Behavioral Data

In order to investigate potential effects of the different training paradigms on performance, absolute error was taken as the main measure of localization accuracy, in addition to the percentage of correct responses. The rationale for using this measure and the computation of absolute error has been described previously in detail (Lewald, 2016). In short, the participants’ responses were assigned to the azimuth indicated by the position of the response button (−60°; −20°; 20°; 60°), and the unsigned deviation of the response from the actual target azimuth was taken as absolute error. Responses to targets presented at ±60° were excluded from analyses since these data did not provide information on errors to more eccentric positions (Lewald, 2016). Absolute errors were normalized with reference to the pre-training block in the same way as described below for ERP data.

### EEG Recording and ERP Analysis

The continuous EEG was sampled at 1 kHz using a QuickAmp-72 amplifier (Brain Products, Gilching, Germany) and 58 Ag/AgCl electrodes, with electrode positions based on the International 10-10 system. Horizontal and vertical electro-oculograms were recorded from four additional electrodes positioned around the left and right eyes. The ground electrode was placed on the center of the forehead, just above the nasion. Two additional electrodes were placed on the left and right mastoids. Electrode impedance was kept below 5 k⋂. The raw data were band pass filtered off-line (cut-off frequencies 0.5 and 25 Hz; slope 48 dB/octave), re-referenced to the average of 58 channels (56 EEG and 2 mastoid electrodes), and segmented into 2000-ms stimulus-locked epochs covering the period from −200 to 1800 ms relative to sound onset. Data were corrected for ocular artifacts using the procedure proposed by Gratton et al. (1983). Individual epochs exceeding a maximum-minimum difference of 200 μV were excluded from further analysis, using the automatic artifact rejection implemented in the BrainVision Analyzer software (Version 2.0; Brain Products, Gilching, Germany). The remaining epochs were baseline corrected to a 200-ms pre-stimulus window and averaged for each subject, separately for each training condition (audiovisual congruency; visual feedback; control) and each test block (pre-training; post-training; 1-h post-training).

Peaks of four primary ERP components (P1, N1, P2, N2) were defined as the maximum positivity or negativity within particular latency windows of specific waveforms after sound onset (P1: 10-110 ms at FCz; N1: 60-160 ms at Cz; P2: 155-255 ms at FCz; N2: 240-340 ms at Cz). The choice of electrode positions was based on previous knowledge of the ERPs topographical scalp distribution (e.g., Anderer et al., 1996; Folstein & Van Petten, 2008; Martin et al., 2008) and confirmed by visual inspection of the grand average waveforms. As we were interested in effects of training on successful sound localization, only trials with correct responses were included in ERP analyses. To take account of placebo and learning effects, ERP data (peak amplitudes and latencies) were normalized by subtraction of pre-training results (baseline) using the formulae:

pre-normalized post-training value = post-training value – pre-training value and

pre-normalized 1-h post-training value = 1-h post-training value – pre-training value.

The pre-normalized data were submitted to 3 × 2 × 2 analyses of variance (ANOVAs) with training condition (audiovisual congruency; visual feedback; control) and block (posttraining; 1-h post-training) as within-subjects factors and group (younger; older) as between-subjects factor. In addition, to investigate effects of training on ERP amplitudes and latencies, pre-normalized data were analysed using one-sample *t*-tests against zero.

### Cortical Source Localization

The cortical sources of the effect of training on ERP amplitudes were localized using standardized low-resolution brain electromagnetic tomography (sLORETA; Pascual-Marqui, 2002), which is part of the LORETA-KEY software package (v20171101) of the KEY Institute for Brain-Mind Research, Zurich, Switzerland (available at www.uzh.ch/keyinst/loreta). Data were baseline corrected to a 200-ms pre-stimulus window for each subject, separately for each training and each test block. Data obtained in the pre-training block were subtracted from data obtained in post-training blocks, and the resulting pre-normalized data for training conditions were contrasted against the related pre-normalized data for the control condition (paired groups, test [A-A2] = [B-B2], with A = post/training, A2 = pre/training, B = post/control, B2 = pre/control). We employed sLORETA within 5-ms time windows around the individual N2 peak-amplitude values (at Cz) for each subject, with the individual N2 latencies taken from the ERP analyses described above.

## Results

### Behavioral data

Both groups showed high levels of percentages of correct responses, differing significantly from chance-level (25%; younger group: *t*[20] = 28.55, *p* < 0.001; older group: *t*[23] = 13.09,*p* < 0.001). For pre-training trials, an ANOVA including the within-subjects factor condition (audiovisual-congruency; visual feedback; control) and the between-subjects factor group (younger; older) revealed a significant main effect of group (*F*[1,43] = 10.30, *p* = 0.003, *η_p_^2^* = 0.19), indicating a higher percentage of correct responses in younger *(M* = 84.3%, SE 2.4%), than older (*M* = 69.9%, SE 3.6%), participants. Also, prior to training, absolute errors were significantly smaller in the younger (*M* = 11.7°, SE 3.1°), than in the older (*M*= 26.5°, SE 2.9°, *F*[1,43] = 11.94, *p* = 0.001, *η_p_^2^* = 0.22), group. There were no significant differences between training conditions within groups in the pre-training blocks, neither in the percentages of correct responses (all *F* ≤ 0.60, *p* ≥ 0.55), nor in absolute errors (all *F* ≤ 0.70, *p* ≥ 0.50).

A 3 × 2 × 2 ANOVA on pre-normalized absolute errors with training condition (audiovisual congruency; visual feedback; control) and block (post-training; 1-h post-training) as within-subjects factors and group (younger; older) as between-subjects factor did not indicate main effects or interactions (all *F* ≤ 3.13, *p* ≥ 0.084). However, across conditions, blocks, and groups, pre-normalized absolute errors were significantly below zero (*t*[44] = −3.98, *p* = 0.0003), thus indicating general improvement in accuracy that was independent of the type of training condition.

### ERPs

In both groups, sound onset elicited a prominent response at vertex position Cz, mainly consisting of a positive deflection (P1), a negative deflection (N1), a second positive deflection (P2), and a second negative deflection (N2; Fig. 2). The latter two components were less prominent in amplitude in older, than younger, subjects, which is in line with previous results (e.g., Getzmann, Hanenberg, et al., 2015). Mean latencies (with reference to sound onset) were 116 ms (*SE* 3 ms) for N1, 213 ms (*SE* 3 ms) for P2, and 296 ms (*SE* 5 ms) for N2 components (averaged across groups and conditions). Pre-normalized data (see *EEG Recording and ERP Analysis)* were submitted to 3 × 2 × 2 ANOVAs, with training condition (audiovisual congruency; visual feedback; control) and block (post-training; 1-h post-training) as within-subject factors and group (younger; older) as between-subjects factor.

**Fig. 2.**
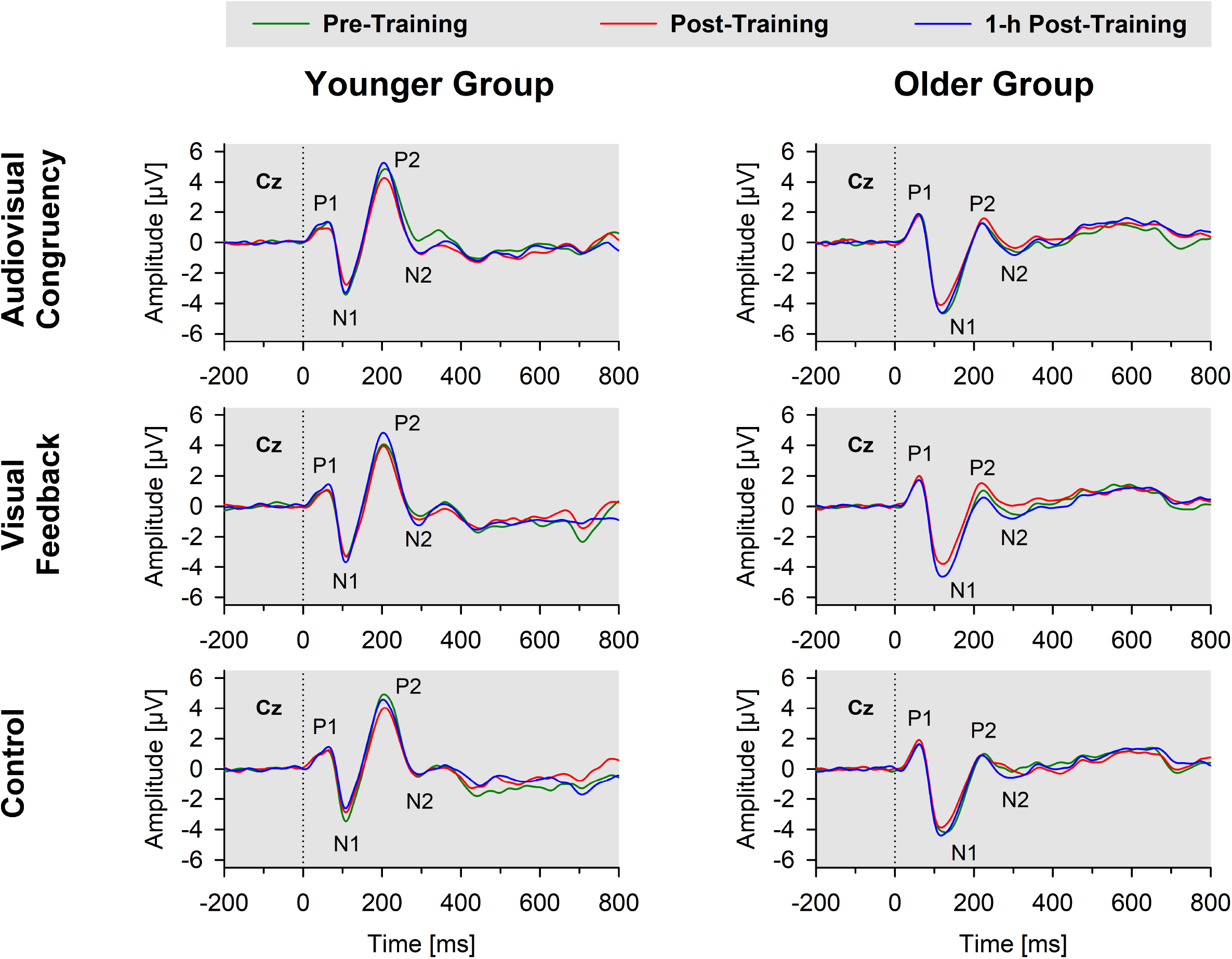
Grand-average ERPs to stimulus onset for younger (left panels) and older subjects (right panels) at Cz electrode position. Data obtained before training (pre-training), immediately after training (post-training), and 1h after training (1-h post-training) are shown separately for audiovisual-congruency training, visual-feedback training, and control condition. P1, N1, P2, and N2 components are marked in each diagram.

### P1 component

For pre-normalized P1 peak amplitudes at electrode position Cz, the ANOVA indicated a significant interaction of condition and block (*F*[1,43] = 3.19, *p* = 0.046, *η_p_^2^* = 0.07), but no further main effects or interactions (all *F* ≤ 3.00,*p* ≥ 0.99). However, *t*-tests against zero, conducted separately for training conditions and blocks, did not reveal significant results (all *t*[44] ≤ 1.33, *p* ≥ 0.19; one-sample *t*-test, two-tailed; Bonferroni-adjusted *α* = 0.008). Also, paired post-hoc *t*-tests, comparing pre-normalized peaks obtained after audiovisual-congruency training or visual-feedback training with the control condition did not reveal any significant differences (all *t*[44] ≤ −1.61, *p* ≥ 0.12; Bonferroni-adjusted α = 0.007). For pre-normalized P1 peak latencies, there were no significant interactions or main effects (all *F* ≤ 1.17, *p* ≥ 0.29).

### N1 component

For pre-normalized N1 peak amplitudes at electrode position Cz, the ANOVA indicated a main effect of block (*F*[1,43] = 9.90, *p* = 0.003, *η_p_^2^* = 0.19), but no further main effects or interactions (all *F* ≤ 1.73, *p* ≥ 0.18). Pre-normalized amplitudes in post-training *(M* = 0.49 μV, *SE* = 0.17; *t*[44] = 2.84, *p* = 0.007), but not in 1-h post-training, blocks (*M* = 0.08 μV, *SE* = 0.20; *t*[44] = 0.39, *p* = 0.70) were significantly in the positive range (one-sample *t*-tests, two-tailed; Bonferroni-adjusted *α* = 0.025), thus indicating reduction in N1 amplitude immediately after training. Pre-normalized peak amplitudes were more positive in posttraining blocks than in 1-h post-training blocks (*t*[44] = 3.21, *p* = 0.002; paired *t*-test, twotailed).

For pre-normalized N1 peak latencies, the ANOVA indicated a significant block × group interaction (F[1,43] = 6.99, *p* = 0.011, *η_p_^2^* = 0.14), as well as a main effect of group (F[1,43] = 7.45, *p* = 0.009, *η_p_^2^* = 0.15). No further main effects or interactions were obtained (all *F* ≤ 1.10, *p* ≥ 0.34). N1 peak latencies significantly differed between groups in the posttraining (younger group: *M* = 3.49 ms, *SE* = 1.56 ms; older group: *M* = −5.26 ms, *SE* = 1.97; *t*[43] = −3.42, *p* = 0.001), but not in the 1-h post-training, block (younger group: *M* = 0.00, *SE* = 1.42; older group: *M* = −2.36 ms, *SE* = 1.60 ms; *t*[43] = −1.09, *p* = 0.28; Bonferroni-adjusted *α* = 0.025). However, *t*-tests against zero for pre-normalized N1 latencies (conducted separately for blocks and groups) did not reveal significant results (younger group: all *t*[20] ≤ 2.24,*p* ≥ 0.04; older group: all *t*[23] ≤ −2.67,*p* ≥ 0.014; Bonferroni-adjusted *α* = 0.0125).

### P2 component

For pre-normalized P2 peak amplitudes at electrode position FCz, there was a significant interaction of block × group (F[1,43] = 9.24, *p* = 0.004, *η_p_^2^* = 0.18), but no further main effects or interactions (all *F* ≤ 2.54, *p* ≥ 0.12). As revealed by post-hoc testing, amplitudes in the younger, but not in the older, group were significantly lower in post-training, than in 1-h post-training, blocks (younger group: post-training *M* = −0.73 μV, *SE* = 0.41 μV, 1-h post-training *M* = 0.24 μV, *SE* = 0.36 μV, *t*[20] = −3.27, *p* = 0.004; older group: post-training *M* = 0.34 μV, *SE* = 0.29 μV, 1-h post-training *M* = 0.04 μV, *SE* = 0.28 μV, *t*[23] ≤ 1.03, *p* > 0.31; two paired *t*-tests, one for each group, two-tailed; Bonferroni-adjusted α = 0.0125). However, *t*-tests against zero for pre-normalized P2 peak amplitudes showed neither significant results when conducted separately for each block (all *t*[44] ≤ −0.62,*p* ≥ 0.54; one-sample *t*-test, two-tailed; Bonferroni-adjusted α = 0.025), nor when conducted separately for each group (younger group: all *t*[20] ≤ −1.77, *p* > 0.09; older group: all *t*[23] ≤ 1.17, *p* > 0.26; one-sample *t*-tests, two-tailed; Bonferroni-adjusted *α* = 0.0125).

For pre-normalized P2 peak latencies, the ANOVA indicated a significant interaction of training condition × group (*F*[*1*,43] = 5.70, *p* = 0.005, *η_p_^2^* = 0.12), as well as a main effect of block (*F*[1,43] = 7.44, *p* = 0.009, *η_p_^2^* = 0.15), with no further main effects or interactions (all *F* ≤ 1.24, *p* ≥ 0.27). Post-hoc testing revealed that pre-normalized peak latencies were significantly shorter than zero in the 1-h post-training block *(M* = −4.38 ms, *SE* = 1.11 ms; *t*[44] = −3.94,*p* < 0.0001), but not in the post-training block (*M* = −0.52 ms, *SE* = 1.44 ms; *t*[44] = −0.36, *p* = 0.72; one-sample *t*-test, two-tailed; Bonferroni-adjusted α = 0.025). Prenormalized latencies, analyzed separately for training conditions and groups, were significantly shorter than zero in the younger, but not in the older, group for the audiovisual-congruency condition in the1-h post-training block, but not for the other conditions and the post-training block (younger group: *M* = −10.19 ms, SE = 3.08 ms, *t*[20] = – 3.31,*p* = 0.003; all other *t*[20] ≤ −2.67,*p* > 0.015; older group: all *t*[23] ≤ −3.11,*p* ≥ 0.005; one-sample *t*-tests, two-tailed; Bonferroni-adjusted *α* = 0.004).

### N2 component

For pre-normalized N2 peak amplitudes at electrode position Cz, the ANOVA indicated a training condition × group interaction (*F*[2,86] = 3.34, *p* = 0.04, *η_p_^2^* = 0.07; Fig. 3), but merely non-significant numerical trends for the factors block (*F*1,43 = 3.70, *p* = 0.06, ¾^2^ = 0.08) and group (*F*[1,43] = 3.79, *p* = 0.06, *η_p_^2^* = 0.08). No further main effects or interactions were found (all *F* ≤ 2.78, *p* ≥ 0.1). Averaged across post-training and 1-h posttraining blocks, pre-normalized peak amplitudes were significantly larger than zero for the younger subjects after audiovisual-congruency training (*t*[20] = −3.39, *p* = 0.003; two one-sample *t*-tests, two-tailed; Bonferroni-adjusted *α* = 0.008), but not for the other conditions (all |*t*[20]| ≤ −1.24, *p* ≥ 0.23). Neither, there were any significant results for the older group with all conditions (all *t*[23] ≤ 1.27, *p* ≥ 0.22; one-sample *t*-test, two-tailed; Bonferroni-corrected *α* = 0.025). In line with that, pre-normalized N2 peak amplitudes in the audiovisual-congruency condition were significantly larger than in the visual-feedback training and control conditions exclusively for the younger group and for the post-training block, (*M* = −1.12 μV, *SE* = 0.34 μV; *t*[20] = −3.29,*p* = 0.004; all other |*t*[20]| ≤ −0.77,*p* ≥ 0.45; one-sample *t*-test, two-tailed; Bonferroni-adjusted α = 0.008), indicating a specific increase in N2 amplitude after audiovisual-congruency training. In the 1-h post-training block, prenormalized N2 peak amplitudes in the audiovisual-congruency condition were numerically larger than in the visual-feedback training and control conditions, again, exclusively for younger subjects (*M* = −1.05 μV, *SE* = 0.36 μV; *t*[20] = −2.90, *p* = 0.009; all other |*t*[20]| ≤ – 1.62, *p* ≥ 0.12; one-sample *t*-test, two-tailed; Bonferroni-adjusted α = 0.008). Post-hoc testing did not reveal any significant differences in the older group (all |*t*[23]| ≤ −2.00,*p* ≥ 0.06; one-sample *t*-test, two-tailed; Bonferroni-adjusted α = 0.008). In line with these findings on N2 peak amplitudes, pre-normalized topographies showed a fronto-central/parietal negativity after audiovisual-congruency training exclusively in the younger group, with its maximum in the left hemisphere (Fig. 4).

**Fig. 3.**
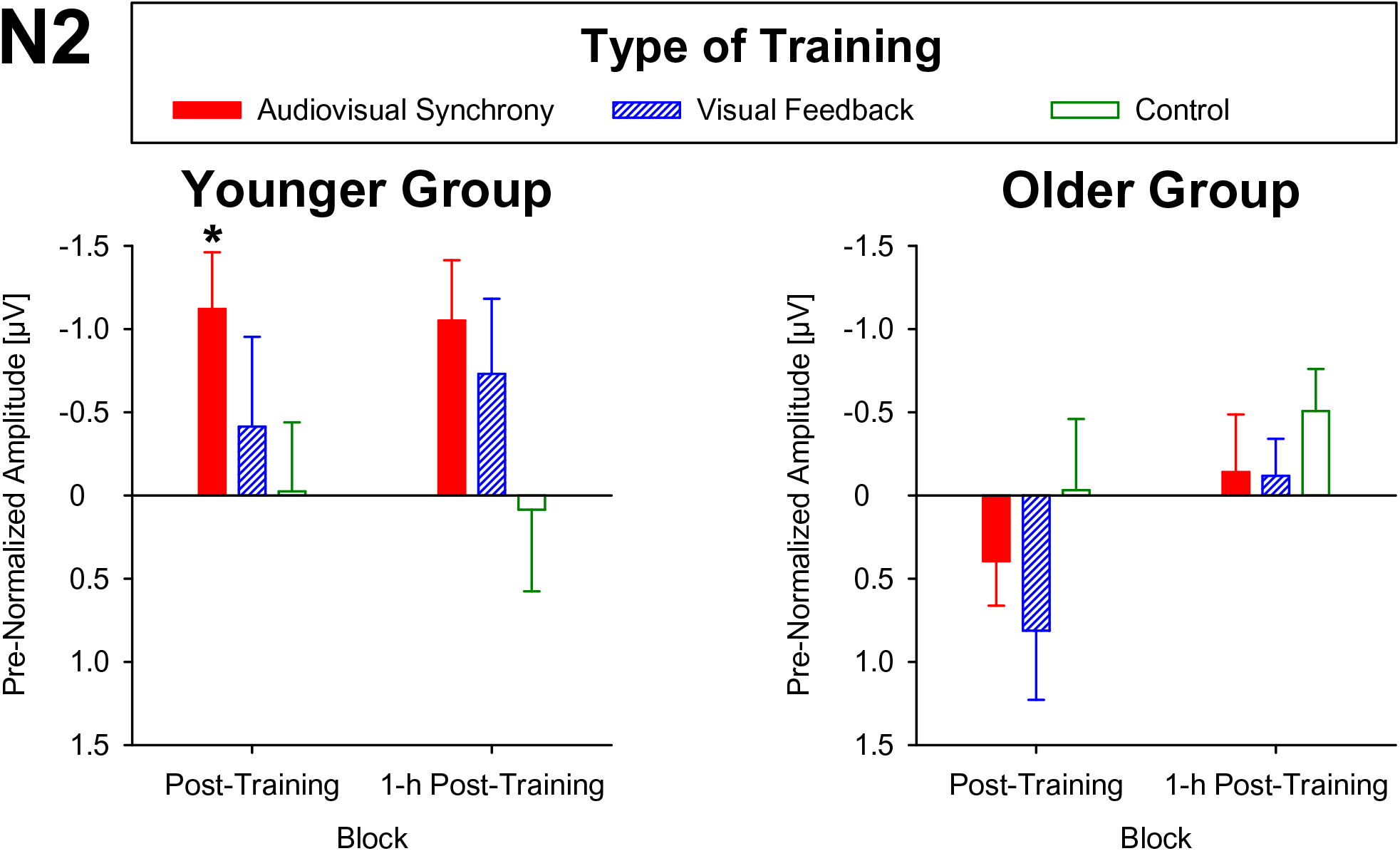
Pre-normalized N2 peak amplitudes at electrode position Cz in younger and older subjects. Post-training and 1-h post-training data are shown separately for audiovisual-congruency training, visual-feedback training, and control condition. A significant negative deviation from zero was found exclusively in the post-training block after audiovisual congruency training in the younger group.

**Fig. 4.**
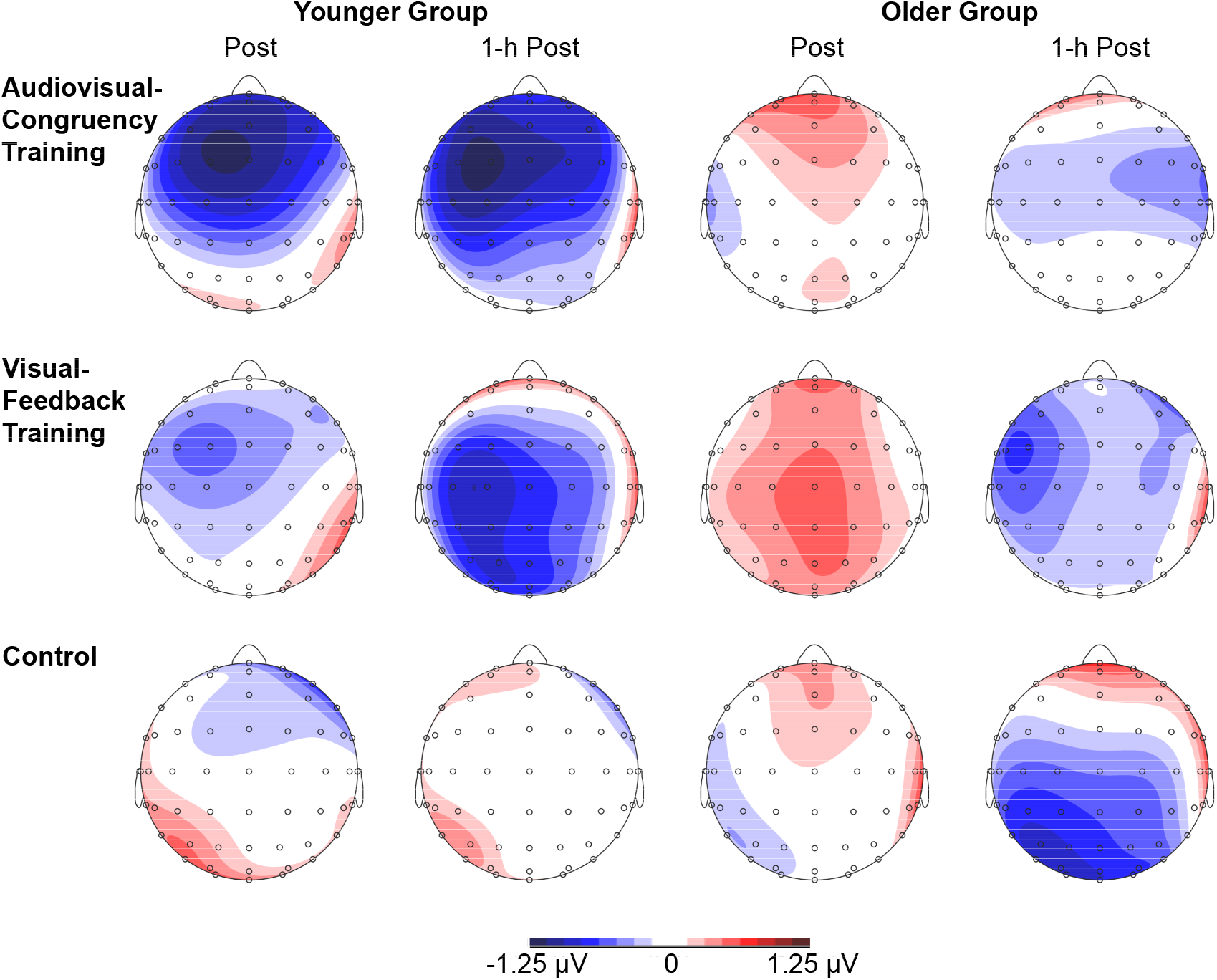
Topographies of N2 components. Pre-normalized topographies, assessed immediately after (post-training) and 1h after training (1-h-post-training) in younger and older subjects are shown separately for audiovisual-congruency training, visual-feedback training, and control condition.

For pre-normalized peak latencies, the ANOVA revealed a significant main effect of block (F[1,43] = 7.10, *p* = 0.011, *η_p_^2^* = 0.14). No further main effects or interactions were obtained (all *F* ≤ 2.09, *p* ≥ 0.13). Averaged across training conditions and groups, N2 latencies were significantly longer in post-training (*M* = 6.28 ms, *SE* = 3.20 ms), than 1-h post-training, blocks (*M* = −0.06 ms, *SE* = 2.27 ms; *t*[44] = 6.34, *p* = 0.009). However, there were no significant differences from zero (all *t*[44] ≤ 1.96, *p* ≥ 0.056; one-sample *t*-test, twotailed; Bonferroni-corrected *α* = 0.025).

### Cortical source of electrical activity at the time of the N2

The cortical source of the enhancing effect of training on the N2 amplitude found for the younger group in the post-training block of the audiovisual-congruency condition was localized using sLORETA. Data obtained in the pre-block were subtracted from data obtained in the post-block and the resulting difference values were contrasted against the related difference values for the control condition. The analysis revealed an enhancement of electrical activity induced by audiovisual-congruency training at a focal peak location at MNI coordinates *X* = 25 mm, *Y* = 50 mm, *Z* = 40 mm (*t* = 6.45, *p* = 0.005, two-tailed), in right superior frontal gyrus (Brodmann area, BA 9) in the anterior region of superior frontal sulcus (SFS; Fig. 5).

**Fig. 5.**
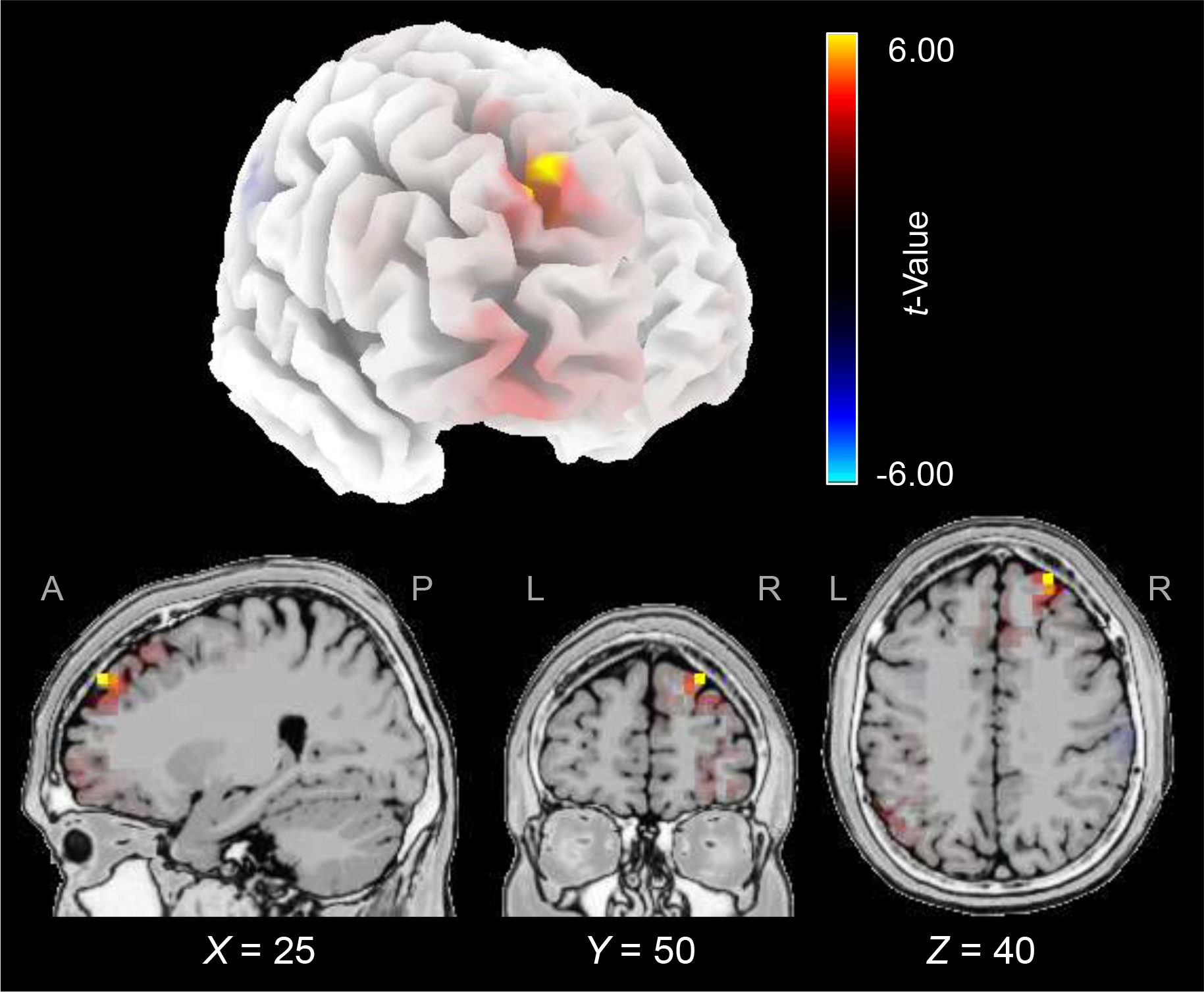
Electrical brain activity at the time of the N2 after audiovisual-congruency training in the younger group. Electrical activity as revealed by sLORETA in the post-block was normalized with reference to pre-training activity and the resulting difference values were contrasted against the related difference values for the control condition. Data were mapped onto a standard 3-D brain template (top) or onto sagittal, coronal, and horizontal slices (T2 MNI-template ‘Colin 27 of sLORETA) positioned at X, Z, and Y coordinates (MNI) as given in the figure (bottom). Color coding shows *t*-values, with warm colors indicating decrease of electrical activity and cold colors indicating increase of electrical activity after training intervention (peak activity: *t* = 6.45, *p* = 0.005).

## Discussion

These results demonstrated an age-specific effect of audiovisual spatial training on neurophysiological correlates of auditory selective spatial attention in a simulated, cocktail-party” scenario. After audiovisual-congruency training, but not after visual-feedback training and the control condition, the N2 peak amplitude was increased. This effect was observed in younger, but not older, participants with a non-significant decline within 1 h after training. At the time of the N2, electrical imaging revealed an increase in activity after audiovisual-congruency training that was located in right dorsolateral prefrontal cortex (DLPFC, BA 9), in the anterior region of SFS. These findings provided, on the one hand, clear evidence that specific training characteristics, namely temporal synchrony and spatial congruency of auditory and visual target stimuli, can enhance ERP correlates of auditory spatial attention. On the other hand, we failed to find a related specificity for behavioural performance, which was generally improved after all training and control conditions.

### Enhanced N2 amplitude after audiovisual congruency training

Audiovisual congruency training induced a specific post-intervention enhancement of the N2, rather than P1, N1, and P2 components, for which no consistent changes were observed. This finding is in alignment with previous studies suggesting the N2 to be a decisive correlate of auditory selective spatial attention in “cocktail-party” scenarios (Gamble & Luck, 2011; Lewald & Getzmann, 2015; Getzmann, Hanenberg, et al. 2015, Lewald et al., 2016; Hanenberg et al., 2019). An enhanced N2 was present not only immediately following training, but also in the 1-h-post block, with a non-significant tendency to decrease as a function of time after training. Thus, it seems as if there were longer-lasting modulations in cortical processing after audiovisual congruency training, which might be due to processes of neural plasticity induced by performing multisensory target localization.

An effect of cognitive training on the N2 has not been reported thus far in the context of auditory spatial attention. Previous studies focussing on improvements of working memory reported increases in N2 amplitude following *n*-back training in healthy subjects (Covey et al., 2019) as well as in Multiple Sclerosis patients (Covey et al., 2018). Thus, on the basis of the limited data available at present, it remains open whether audiovisual congruency training had an effect on N2-related processes specifically involved in auditory attentional functions or rather enhanced more general cognitive processes associated with the N2.

The N2 enhancement following training was restricted to the younger group, which was an unexpected finding. The absence of an effect of training in the older group could possibly be due to the less flexible cognitive system of older adults, which is generally more occupied by task difficulty than that of younger persons (Gajewski & Falkenstein, 2012; Getzmann, Falkenstein,et al., 2015; Getzmann, Wascher, et al., 2015, Getzmann, Hanenberg, et al., 2015; Olfers & Band, 2017; Salthouse, 2009; Yordanova et al., 2004). Our finding may be compatible with ERP research on age differences in speech perception. While older adults often showed reduced N2 amplitudes (Daffner et al., 2015), probably reflecting a less successful reorienting toward the stimulus of interest, younger adults showed more pronounced N2 amplitudes, reflecting an enhanced inhibitory control (Getzmann, et al., 2014; Getzmann, Wascher, et al., 2015). Thus, that the N2 enhancement was observed here in younger, but not older, adults could indicate a specific impact of the audiovisual-congruency training on processes of cognitive control. The question of whether related ERP enhancements, as found here for the N2 in younger participants, can also be induced in elderly people by more intense and repeated daily training over longer periods has to be addressed by future studies.

### Cortical sources of training-induced N2 enhancement

At the time of the N2, an increase of electrical activity after audiovisual-congruency training was found to be located in right superior frontal gyrus, in the anterior region of SFS (cf Fig. 5). Previous studies, focussing on the cortical correlates of selective auditory spatial attention using various methods, have revealed several cortical regions, composing a complex network. This network comprises auditory cortex, posterior superior temporal gyrus (pSTG) and planum temporale (PT), inferior parietal lobule (IPL), superior parietal lobule and precuneus, inferior frontal gyrus, frontal eye field (FEF), as well as regions of BA 9 and SFS, which were nearby the location of training-induced activity change found here (Pugh et al., 1996; Nakai, Kato, & Matsuo, 2005; Braga et al., 2016, 2017; Lee et al., 2014; Lewald, 2016, 2019; Lewald & Getzmann, 2015; Lewald et al., 2016, 2018; Kong et al., 2014; Zündorf et al., 2013, 2014). In particular, the area including SFS and FEF has been related to the so-called N2ac (Lewald et al., 2016), an anterior contralateral N2 subcomponent, which has been regarded as a correlate of auditory selective spatial attention (Gamble and Luck, 2011). Generally, the SFS is well-known as auditory spatial region of dorsofrontal cortex. This region has been demonstrated in many studies to be involved in sound localization (e.g., Alain et al., 2001; Weeks et al., 2000; Zatorre et al., 2002; Lewald et al., 2008; Zündorf et al., 2016) and has been assigned to the auditory posterodorsal (spatial) pathway (Arnott et al., 2004). However, it has to be noted that the activity increase revealed here was located at a more anterior position of SFS, compared with areas described in the studies cited above, which reported positions in caudal SFS. Because of the low spatial resolution of the electrical imaging method used here, any clear-cut conclusions on the location of activity in specific subareas of DLPFC might be difficult to draw, and further studies using imaging techniques with higher spatial resolution, such as functional magnetic resonance imaging (fMRI), may have to clarify this issue.

The present results may be related to recent findings by transcranial direct current stimulation (tDCS). In a preceding study using the same “cocktail-party” task as used here, a significant enhancement of the N2 was observed after monopolar anodal tDCS of right pSTG, including PT and auditory cortex (Hanenberg et al., 2019). Also, bilateral monopolar anodal tDCS over this area has been shown to induce clear offline improvements in behavioural performance with this task (Lewald, 2019). These findings have been related to the crucial role of PT in “cocktail-party” sound localization, as had been revealed by fMRI in healthy persons (Zündorf et al., 2013) and voxel-based lesion-behaviour mapping analyses in stroke patients (Zündorf et al., 2014). The PT may represent an initial stage of auditory spatial processing within the hierarchically organized posterodorsal cortical pathway, channelling information to frontoparietal areas for further analyses (Griffiths & Warren, 2002; Krumbholz et al., 2005), including those relevant for selective spatial attention (Zündorf et al., 2014). In general alignment with this view, Hanenberg et al. (2019) found a reduction of activity in right IPL at the time of the enhanced N2 after anodal tDCS over right pSTG. The IPL is connected with ipsilateral DLPFC via dorsal components of the superior longitudinal fasciculus (SLF), a white-matter bundle that is crucially involved in functions of spatial orienting and awareness, as well as attentional control (Bernal & Altman, 2010; Doricchi & Tomaiuolo, 2003; Makris et al., 2005; Thiebaut de Schotten et al., 2005, 2008; Doricchi et al., 2008; Suchan et al., 2014; Nakajima et al., 2020). Zündorf et al. (2014) showed that left-sided lesions of the SLF were associated with deficits in the “cocktail-party” task, suggesting an important role of this structure and its frontal target areas in auditory selective spatial attention. Thus, the DLPFC might be part of a temporo-parieto-frontal network concerned with auditory functions subserving “cocktail-party” listening.

The specific post-intervention increase in activity found after audiovisual-congruency training suggests that the related processes in DLPFC were specifically strengthened by bimodal stimulation. This result may be compatible with previous findings indicating multi-or supramodal properties of the dorsofrontal networks that have been usually associated with selective spatial attention in the visual modality (e.g., Bharadwaj et al., 2014; Lee et al., 2012; Lewald et al., 2018; Macaluso, 2010; Slotnick & Moo, 2006). For the monkey DLPFC, it has been suggested that neuronal processes exist for visual and auditory location information and spatial working memory (Fuster et al., 2000; Kikuchi-Yorioka & Sawaguchi, 2000; Artchakov et al., 2007; Hwang & Romanski, 2015; for review, see Plakke & Romanski, 2016), and the human DLPFC has been shown to be involved in transforming auditory and visual inputs into multimodal spatial representations that can be used to guide saccades (Tark & Curtis, 2013). The monkey DLPFC receives projections from posterior auditory cortex areas known to be involved in spatial processing and from the posterior parietal cortex (Chavis, & Pandya, 1976; Rauschecker, et al., 1995; Romanski, Bates, et al., 1999; Romanski, Tian, et al., 1999). The latter area, which also has reciprocal connections with posterior auditory cortex, has been shown to be critically concerned with auditory and visual spatial processing in human and nonhuman primates (e.g., Mishkin et al., 1983; Bushara et al., 1999; Mazzoni et al., 1996; Romanski, Tian, et al., 1999; Lewald et al., 2002, 2004, 2016). Thus, in conclusion, we assume that the present finding of activity enhancement in BA 9 induced by repetitive processing of spatially and temporally congruent audiovisual stimuli during training may be related to the auditory-visual bimodal properties of the dorsal attention network composed of the DLPFC region and its connections with posterior parietal areas via SLF.

### Audiovisual congruency as a key factor for training effects

An effect of training on the N2 was found exclusively for the audiovisual-congruency condition. This result may be in alignment with the multitude of studies on audiovisual integration, which have demonstrated bimodal enhancement by spatiotemporal alignment using several behavioral (Alais & Burr, 2004; Lewald, 2002b; Lewald & Guski, 2003; Lewald et al., 2001; Lovelace, et al., 2003; Pages & Groh, 2013; Recanzone, 1998) and neurophysiological approaches (Besle et al., 2004; Santangelo et al., 2008; Stein & Stanford, 2008; Stekelenburg & Vroomen, 2007, 2012; Talsma et al.,2007; for review, see Stein & Meredith, 1993). Also, few studies in animals and humans have already demonstrated at the behavioral level that spatiotemporally congruent audiovisual stimulation can be used to improve accuracy of localization of single sound sources (Cai et al., 2018; Isaiah et al., 2014; Kumpik et al., 2019; Strelnikov et al., 2011; Berger et al., 2018). Even though we failed to find specific effects of audiovisual-congruency training at the behavioral level, the present study extended these previous approaches by showing that electrophysiological correlates of audiospatial attention in the presence of multiple distractor sources were enhanced by this type of training, while no effect was observed for visual-feedback training.

We assume that this result is related to the experience of phenomenal causality of auditory and visual events (i.e., the impression of a common cause) during audiovisual-congruency training, as typically occurs in the ventriloquism effect (Lewald & Guski, 2003). It is important to note that such binding phenomena do not require complex stimuli, with a highly compelling, meaningful association of auditory and visual information. Rather, simple light spots and tone bursts have been shown to be sufficient to induce audiovisual binding if presented in close spatiotemporal proximity (e.g., Thomas, 1941; Bertelson & Radeau, 1981; Radeau & Bertelson, 1987; Lewald et al., 2001; Slutsky & Recanzone, 2001; Lewald & Guski, 2003). In the audiovisual-congruency training used here, light flashes and target animal vocalizations were presented with identical stimulus onset and duration from roughly the same location. Participants may have experienced binding of auditory and visual events, if distractors were successfully suppressed by the occurrence of the “cocktail-party” effect. That is, this type of training may have induced neural processes resulting in more effective distractor suppression and, thus, more accurate representation of auditory targets in the presence of distractor sources. These processes can be described in terms of short-term neural plasticity based on the ventriloquisms effect, as has been discussed in the context of the normally occurring continuous calibration of auditory space by visual experience or its counterpart, the ventriloquism after-effect, in which repetitive or trial-wise presentation of synchronized auditory and visual stimuli with consistent spatial disparity shifts the representation of auditory space relative to the visual space (Recanzone, 1998; Lewald, 2002b). In an EEG source-imaging study focussed on the neural basis of the ventriloquism aftereffect, Park and Kayser (2020) recently reported that prolonged exposure to consistent auditory-visual discrepancies recruits, in addition to sensory (occiptal and temporal) cortices and multisensory parietal areas, prefrontal regions, including inferior frontal, middle frontal, and superior frontal gyri about 240 ms after stimulus onset. This finding could, potentially, be related to the enhancement of DLPFC activity found here after audiovisual-congruency training at the time of the N2. Taken together, spatiotemporal congruency of auditory and visual stimuli during training appears to be a key feature enhancing neural processes of auditory selective spatial attention. Our results suggested that this training-induced short-term plasticity occurs particularly in the DLPFC region at the time of the N2 component of the ERP.

### Training-induced effects on N1 amplitude and P2 latency

Only minor and rather nonspecific post-intervention changes were observed for N1 and P2 components, and no consistent effects at all for the P1 component. The N1 amplitude was generally reduced in the post-training-blocks compared with 1-h-post blocks, independently of the training condition. In terms of learning, changes in N1 amplitude are often referred to as early automatic stimulus processing (Lange, 2013; Näätänen & Picton, 1987) depending on attentional phenomena (Eimer, 2014; Hillyard et al., 1973; for review, see Luck et al., 2000; McEvoy et al., 2001). Increased N1 amplitudes have been associated with attention catching properties of auditory stimuli and task difficulty (Näätänen et al., 2011). On the other hand, decreased N1 amplitudes could indicate less attentional effort after training due to improved early processing of the stimuli, task familiarity, or the participants’ impression that the task was less demanding after already having performed it (Tallus et al., 2015). These factors could also be relevant for our findings, given that the participants performed the test blocks tree times per session.

P2 latencies were specifically shortened in the younger group 1 h after audiovisual-congruency training, but not for the other conditions, the older group, and the post-training block. On the one hand, this result suggests an accelerated occurrence of P2-related processes after this type of training. On the other hand, it seems difficult to interpret since one may generally expect stronger effects of training on ERPs immediately after the interventions, rather than delayed by 1 h. As the analyses also revealed a general shortening of P2 latencies in the 1-h-post blocks, compared with post blocks, it seems likely that an unspecific effect occurred that was due to the repeated execution of the task, independently of the type of intervention. To which extent such unspecific effects have superimposed potential effects specific to the audiovisual-congruency training remains unclear.

### Limitations

Unlike the clear-cut electrophysiological result, task performance was found to be unspecifically improved after all training interventions and independently of age. This outcome was probably due to the multiple repetition of the task, as is often found in research on learning and memory (Ebbinghaus, 1964). That we failed to find specific training effects could, possibly, be a result of ceiling effects (Brungart & Simpson, 2007) after repeated training, as were found previously in a similar task (Hanenberg et al., 2019). However, given mean rates of correct responses of about 84% in the younger and 70% in the older group, ceiling effects should not play a major role here. Alternatively, the effect of the audiovisual-congruency training observed for the N2 amplitude could be confined to specific subprocesses required to solve a “cocktail-party” speech localization task. Assuming that the N2 especially reflects cognitive control processes mainly related to the inhibition of task-relevant information (as argued above), the audiovisual congruency training might have enhanced specific cognitive control processes (reflected by the increase in N2 amplitude), rather than speech-in-noise localization in general.

This study left open the question of whether more extended and more long-lasting improvements in task performance in “cocktail-party” scenarios could be achieved by training in both, younger and older adults. Motivation has been shown to have a significant impact on task performance (Green & Seitz, 2015), and future studies offering more attractive training paradigms, such as game-based training, should be considered here. Recent work on this topic has shown that various age groups profit from action video game training, showing enhanced performances in task switching abilities after playing for three weeks (Basak et al., 2008; Strobach et al., 2012; Wang et al., 2016). Recently, Schuchert and Lewald (2020), using a similar “cocktail-party” task as used here, demonstrated that both audio action game training and video non-action game training improved auditory selective spatial attention in younger adults. The present results suggest that a bimodal (audio-visual synchronous) game training may also be promising in this respect.

### Conclusion

In summary, the present study showed that short-term audiovisual-congruency training, but not visual-feedback training and a control condition, enhanced the N2 component in a multiple-speaker target localization task. The increase in N2 was associated with an increase of electrical activity in DLPFC and may indicate enhancement of neural processes of auditory selective spatial attention. Both effects were observed in younger, but not older, participants. Further experiments are necessary in order to examine whether more intensive, longer lasting and more realistic audio-visual training settings are suitable to obtain improvements also in behavioral measures.

## Acknowledgements

The authors wish to thank Jonas Heyermann and Stefan Weber for help in running the experiments, and Tobias Blanke and Peter Dillmann for preparing software and parts of the electronic equipment. This work was supported by the German Federal Ministry of Education and Research in the framework of the TRAIN-STIM project (Grant number 01GQ1424E).

## Notes

### Competing Interest Statement

The authors have declared no competing interest.

